# Synthetic stimuli reveal a predictive and switch-like activation of the songbird’s vocal motor program

**DOI:** 10.1101/251256

**Authors:** Alan Bush, Juan F. Döppler, Franz Goller, Gabriel B. Mindlin

## Abstract

Acquisition and maintenance of complex vocal behaviors like human speech and oscine birdsong require continuous auditory feedback. The exact way in which this feedback is integrated into the vocal motor programs is not completely understood. Here we show that in sleeping zebra finches (*Taeniopygia guttata*), the activity of the song system selectively evoked by playbacks of their own song can be detected in the syrinx. Measuring the electrical activity of syringeal muscles, we found playback-evoked patterns identical to those recorded during song execution. Using this global and continuous readout we studied the activation dynamics of the song system elicited by different auditory stimuli. We found that a synthetic version of the bird’s song, rendered by a physical model of the avian phonation apparatus, evoked exactly the same response, albeit with lower efficiency. Analysis of these responses reveal a predictive and switch-like activation of the motor program, with preferred activation instants within the song.

**Significance:** The study of the integration between sensory inputs and motor commands has greatly benefited from the finding that in sleeping oscine birds, playback of their own song evokes highly specific firing patterns in neurons also involved in the production of that song. Nevertheless, the sparse spiking patterns that can be recorded from few single neurons gives limited information of the overall activity of the song system. Here we show that this response is not limited to the central nervous system, but reaches vocal muscles. Combining this integrated measure of the activity of the system with surrogate synthetic songs, we found an all-or-nothing and predictive activation of the system, suggesting the existence of a pre-programmed internal dynamics.

## Introduction

The cooperation between different individuals, or the ability to perform coordinated and even synchronic actions, is at the foundation of many species success. This requires a delicate integration between sensory and motor programs, at specific regions of the nervous system. What is the nature of this integration? Let us analyze this problem in the framework of oscine song production.

Our model will be the elicitation of motor like patterns in the oscine song system when exposed, during sleep, to their own song. The avian auditory pathway indirectly connects to the song system, a set of neural nuclei necessary to generate the motor gestures required for phonation (Figure 1a). Neurons of HVC and RA (two of these nuclei) show a stereotyped sparse spiking pattern elicited by playback of autogenous song in sleeping or anesthetized animals, firing at few specific instants of the song (1–4). This response is highly specific to the bird’s own song (BOS) as it is not evoked by similar conspecific songs or a reversed version of BOS (2). Furthermore, the spiking times are almost identical to those of the same neurons measured during song execution (3). This highly selective response to playbacks provides an excellent experimental opportunity to study the activation of the song system by auditory stimuli, and it will be our tool to explore the integration between the auditory and motor programs.

**Figure 1.**
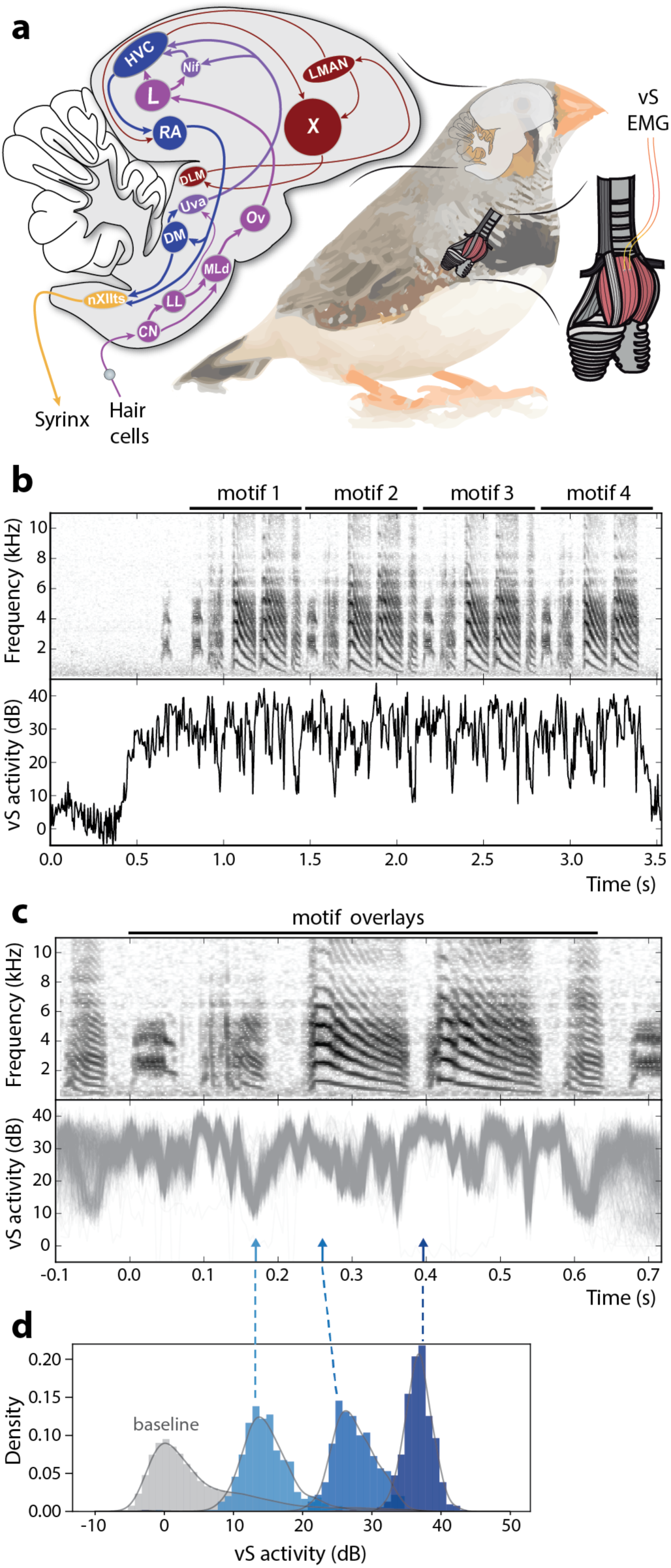
*syringealis ventralis* electromyograms (vS EMG) during song execution. **a)** Top left, schematic representation of the nuclei of the auditory pathway (violet), posterior vocal pathway (blue) and anterior forebrain pathways (red). Bottom right, schematic representation of the syrinx and associated muscles with approximate implantation point of the bipolar electrodes in vS (red). **b)** Top, spectrogram of a typical song of a male zebra finch. Solid lines above the plot indicate song motifs. Bottom, vS activity measured during song execution (bird ab09, 1^st^ surgery). **c)** Top, spectrogram of a single motif. Bottom, overlay of all vS activity traces measured during motif execution. **d)** Distribution of execution vS activity at the indicated times of the motif (blue histograms) and baseline vS activity (gray histogram).

One way to understand how the song system is activated by this stimulus is to study the response to slight variations of BOS that maintain the capacity to activate the system, but in a slightly deteriorated way. In this regard, synthetic songs generated with a mathematical model of the avian vocal organ (SYN) have been shown to evoke firing patterns in HVC similar to those evoked by BOS. Nevertheless, the probability of any given neuron of responding to SYN was at most 60% of that of responding to BOS (5). This suggests that these synthetic songs are ideally suited to study the activation dynamics of the song system by auditory stimuli: good enough to evoke the highly specific response, yet sufficiently different as to produce measurable changes in that response.

But, how do the responses elicited by the BOS and the synthetic stimulus differ? Does SYN produce a partial activation of the song system throughout the song? Or does it produce a full activation but with lower efficiency? The former case would suggest a local response to suboptimal auditory stimuli, while the latter scenario is consistent with the recruitment of a preprogrammed dynamics. In order to test these hypotheses, we need a global, continuous and graded readout of the song system’s activity.

Measuring the sparse firing pattern of few individual neurons gives limited information on the overall activity of the song system. We can overcome this difficulty by measuring efferent motor commands of the system. In this regard, the electrical activity of the *syringealis ventralis* (vS), the largest muscle in the syrinx (Figure 1a), has been measured during song execution (6, 7). Remarkably, this muscle spontaneously activates during sleep, showing variable patterns which sometimes resemble the execution pattern (8).

In this work, we measure electromyograms (EMG) of the vS muscle during song execution and nocturnal auditory playback experiments. This enables the study of the overall activation dynamics of the song system elicited by autogenous and synthetic stimuli with high temporal resolution.

## Results

We successfully implanted electrodes in the syrinx of seven zebra finches, as assessed by the EMG signal acquired. The amplitude of the signal during song execution was in the range of [0.4-1.7] mV before amplification, with a dynamic range around 30 dB compared to baseline fluctuations. Implants lasted from a few days up to three weeks, showing a mild reduction of EMG amplitude over time in some cases. We confirmed electrodes placement in vS by postmortem examinations.

In some animals we observed an electrocardiogram-like (ECG) component in the signal (Figure S1a-b), consistent with the heart rate of small birds (9). The syrinx is a deep structure located close to the heart; therefore, an ECG component in the signal is not surprising. In order to extract the EMG component of the signal we applied the Teager energy operator (10), a measure of local energy of the signal, as we noticed that this operator greatly attenuates the ECG component (Figure S1a). We then estimated the vS activity as the log-ratio of the Teager energy to its baseline value (see details in methods).

Traces of vS activity show a repetitive pattern during song execution, consistent with the reiterations of the song’s motif (Figure 1b). Overlaying these traces for all executions of the motif shows a highly stereotyped pattern, with relatively small variations across renditions (Figure 1c, S1c-d). Furthermore, this measure of vS activity has an approximately normal distribution at each time of the motif (Figure 1d).

In three birds, we performed a second surgery in which we implanted the electrodes in a different position of the vS muscle. The EMG patterns observed in both surgeries were strikingly similar (Pearson’s correlation in [0.53-0.82], p<10^−5^ in all cases, Figure S1e), suggesting a homogeneous activation of the muscle.

Next, we performed nocturnal playback experiments. The auditory stimuli included a recording of the bird’s own song (BOS), a reversed version of the same recording (REV), a conspecific song (CON) and a synthetic version of the bird’s song (SYN), presented in random order (see details in methods). In Figure 2 we show all the vS activity traces measured for a bird, time-locked to each stimulus motif onset. (See Figure S2 for additional birds.) For REV and CON motifs, the traces fluctuate around baseline, with occasional and inconsistent increases in activity. In contrast, a significant fraction of traces in response to BOS show a consistent activation, resulting in an overall pattern reminiscent of the execution activity (Figure 2a). Therefore, playback of the bird’s own song selectively elicits a stereotyped activation in the syrinx, consistent with the response observed in the central nervous system (CNS) (2).

**Figure 2.**
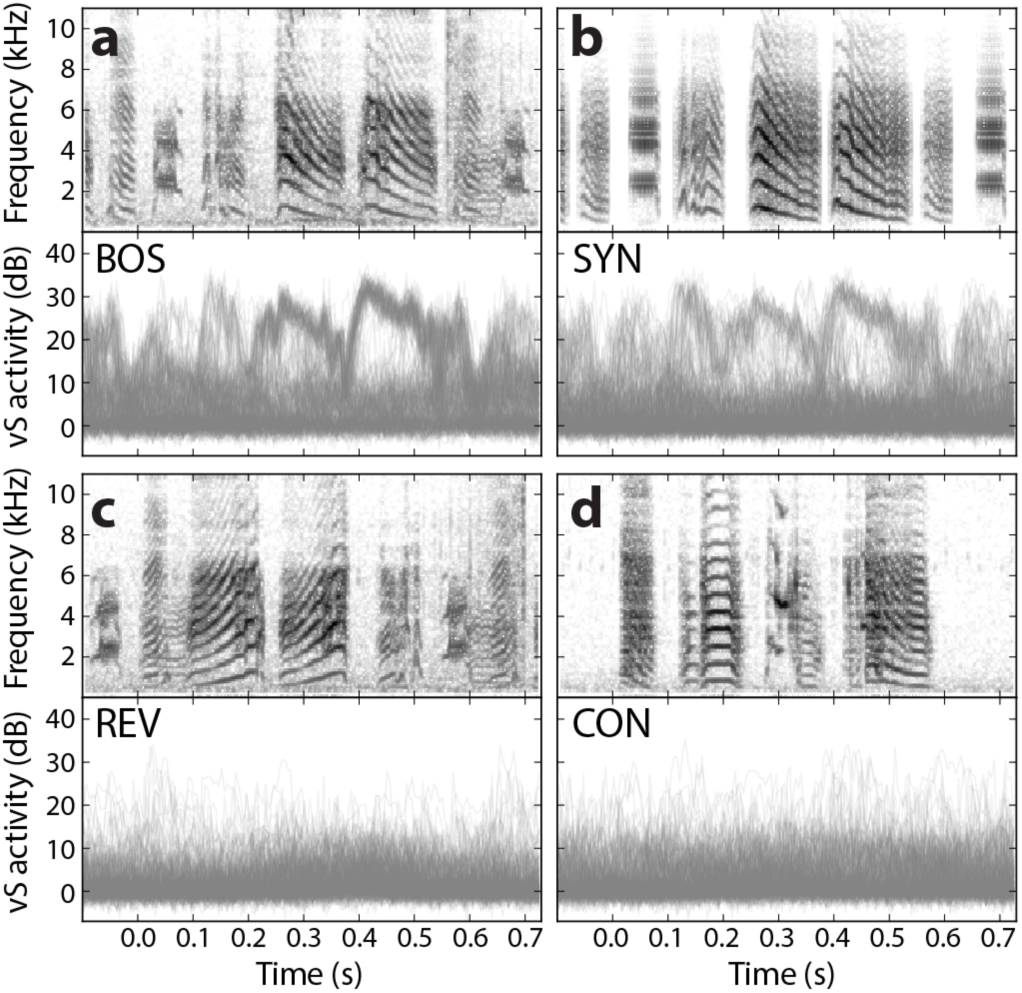
Nocturnal playback responses to the bird’s own song (BOS, **a**), synthetic version of the song (SYN, **b**), reversed version of the song (REV, **c**) and a conspecific song (CON, **d**). In all cases, the top panel shows the spectrogram of the stimulus motif and the bottom panel an overlay of all recorded vS responses to those motifs (bird ab09, 2^nd^ surgery).

Similarly, SYN also stimulated consistent activation of vS. Furthermore, this activation pattern is strikingly similar to that of BOS, although it was evoked in a smaller fraction of the trials (Figure 2b and S2). Hence, our song synthesis is also capable of eliciting selective stereotyped vS activity, consistent with the response observed in HVC (5).

To quantitatively characterize the response to different stimuli we performed a correlation analysis. For each stimulus, we approximated the response patterns as the 95^th^ percentile of vS activity at each time point of the motif (referred to as BOS_95_, SYN_95_, CON_95_ and REV_95_; Figure 3a). We then cross-correlated these patterns to the median vS activity during motif execution (EXE_50_; Figure 3b). BOS_95_ and SYN_95_ had Pearson correlation coefficient ranging from 0.63 to 0.83 (Figure 3c, Table S1), with no significant difference among them (p=0.20), but both were significantly higher than those of CON_95_ and SYN_95_ (p<10^−5^, ANOVA with orthogonal contrasts). The time delays of BOS_95_ and SYN_95_, with respect to the execution pattern EXE_50_, were less than 30ms in all cases, with some cases showing no detectable delay (Figure 3c, Table S1). There was no significant difference in delay times between SYN_95_ and BOS_95_ (p=0.43, paired t-test).

**Figure 3.**
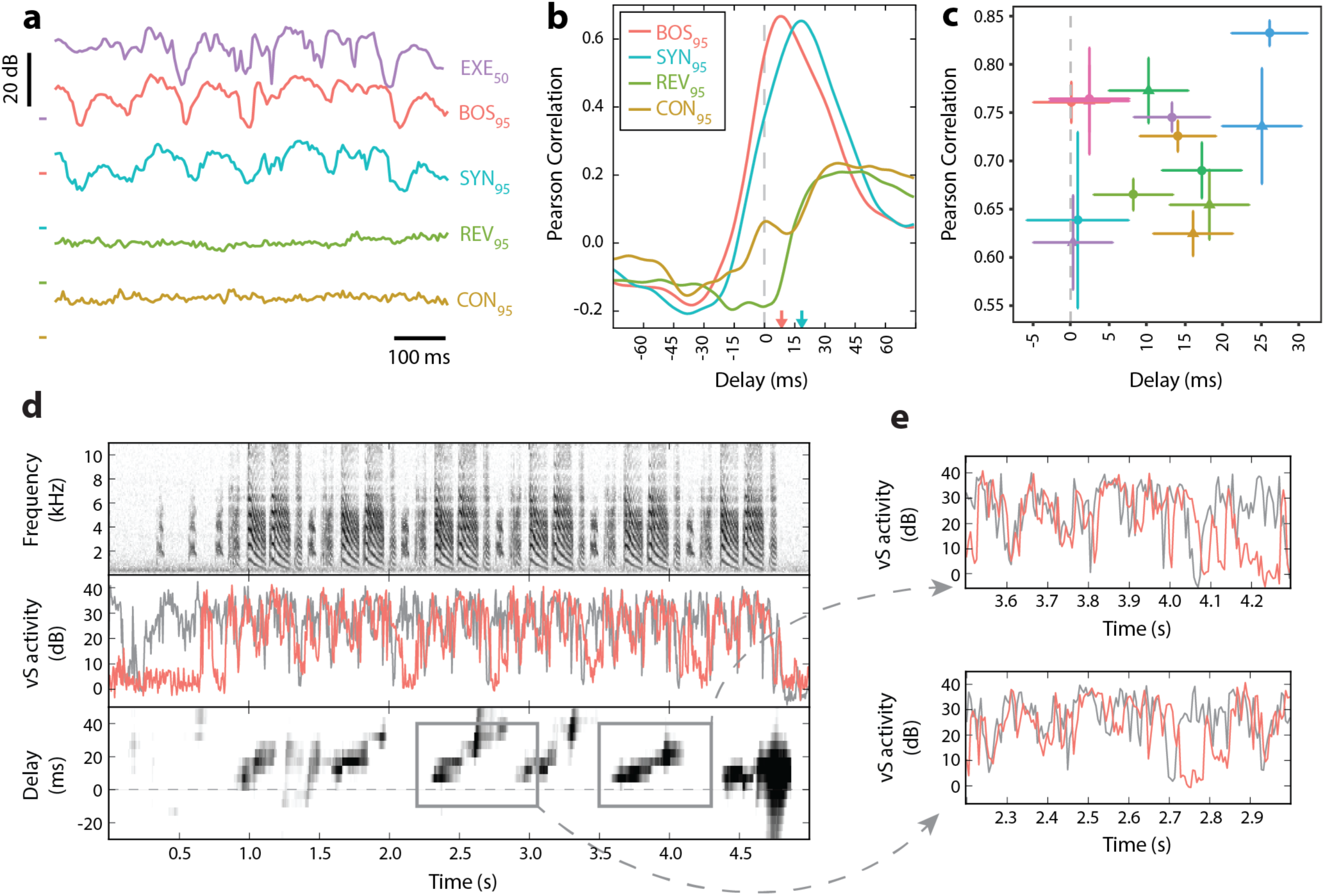
Evoked vS activity correlates with the execution pattern. **a)** Median vS activity pattern during motif execution (EXE_50_) and 95^th^ percentiles of vS activity responses to different auditory stimuli motifs (BOS_95_, SYN_95_, REV_95_ and CON_95_ for bird ab09). **b)** Cross-correlation analyses of responses to auditory stimuli with respect to median execution profile (EXE_50_). Arrows indicate times of maximal correlation for BOS_95_ (red) and SYN_95_ (blue). **c)** Maximal correlation values and correspondent delays for BOS_95_ (circles) and SYN_95_ (triangles) of all animals (see details in Table S1). Horizontal error bars represent uncertainty in the delay calculated by bootstrapping and considering the instrumental error (vS activity was calculated at 200Hz, see methods). Vertical error bars represent the standard error of the maximal correlation coefficient calculated by bootstrapping. **d)** Top, spectrogram of an entire song of bird ab09 used as stimulus. Middle, the execution vS activity profile during execution of that song (gray curve) and the evoked profile during a nocturnal playback (red curve). Bottom, correlation spectrogram showing in gray scale the local (250ms) correlation coefficients between the above traces for every time and delay (see methods). **e)** vS activity for execution (gray) and evoked response to BOS (red) for the indicated time windows.

Upon closer examination, we noticed that for some animals the time delay of individual responses was not constant (Figure 3d-e). To visualize how the delay varies along the motif we constructed “correlation spectrograms”, in which local correlation coefficients to EXE_50_ are shown for all times and delays (bottom panel of Figure 3d, see details in SI Methods). In this particular example, the delay of maximal correlation increases linearly with time, reaching values close to 40ms, after which the correlation is briefly lost and later reappears with a near-zero delay.

To analyze the variability in the playback-elicited vS activity, we time-warped these patterns to a reference execution trace (see SI Methods and Figure S3c). In this way, we could assess the similarity among responses independently of small phase differences between them due to variable time delays.

We performed multidimensional scaling (MDS) analyses on all the vS activity traces measured for the different stimuli and song execution (Figure 4a and S4, see details in SI Methods). MDS is a visualization technique that attempts to represent the multidimensional distance between traces in a 2D plot, *i.e.* proximate points in the MDS plot represent similar vS activity traces. Points representing execution traces (EXE) form a compact cluster towards the right of the plot, while REV and CON points form a disperse cloud towards the left (Figure 4a). Some of the BOS points are in the region of the REV-CON cloud, while others form a cluster towards the top-right of the plot, which marginally overlaps with the execution cluster. Remarkably, SYN points have a similar distribution to that of BOS points, although a smaller fraction of them can be observed in the cluster adjacent to the execution points. The same overall distribution can be observed for other birds (Figure S4).

**Figure 4.**
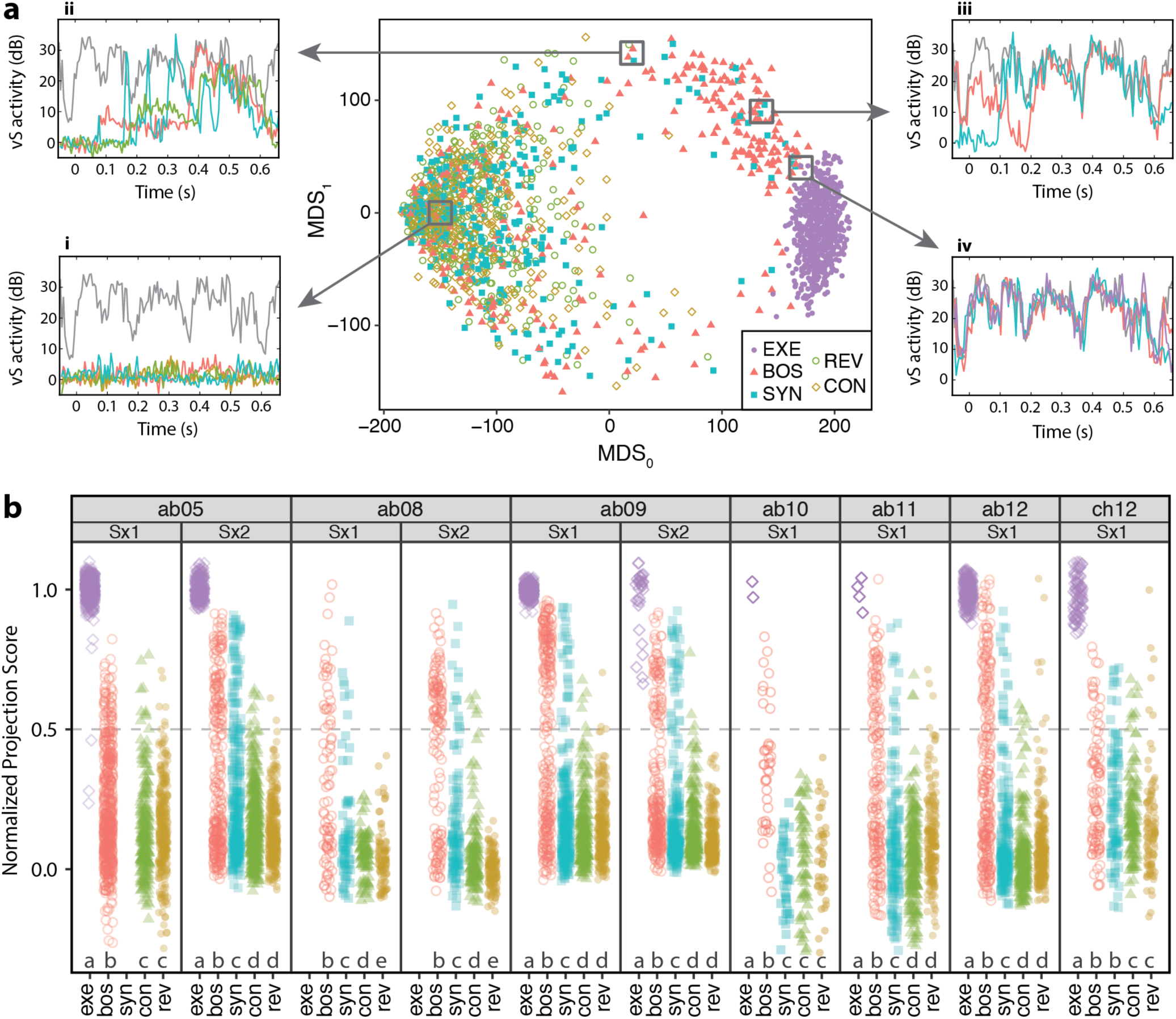
BOS and SYN have similar elicited response distributions. **a)** Multi-Dimensional Scaling (MDS) plot for all execution (EXE, violet) and stimuli-evoked (BOS, red; SYN, blue; REV, green; CON, brown) vS activity traces (bird ab09). Top and right panels, example vS activity traces of the indicated regions of the MDS plot. Gray curve in all panels corresponds to the median execution activity (EXE_50_). **b)** Normalized projection Score (see methods) of all vS activity traces of all animals, surgeries and stimuli. Shared letters (a-d) indicate non-significant differences between conditions (pairwise K-S tests corrected by FDR, see methods). Horizontal dashed line indicates the “high” response threshold. (For ab08, BOS_95_ was used as reference pattern for score calculation instead of EXE_50_ as no execution was recorded.)

Panels i-iv in Figure 4a show example traces corresponding to the indicated points. The median execution profile (EXE_50_) is shown in gray for reference. As can be observed, points in the REV-CON cloud correspond to traces with baseline vS activity (panel i). On the other hand, points towards the right, in the intersection of the BOS-SYN and EXE clusters, correspond to traces that closely follow EXE_50_ (panel iv). Interestingly, other points in the BOS-SYN cluster correspond to traces that closely follow the execution profile, but for only a portion of the motif (panels ii-iii).

In order to perform a statistical analysis, we defined the normalized projection score (*ρ*), a measure of similarity between vS activity traces and median execution profile (see details in methods). A value of zero indicates baseline vS activity, while unity corresponds to a trace exactly matching EXE_50_ in shape and amplitude. The distribution of this score is shown in Figure 4b for all birds and stimuli. As expected, EXE traces have scores close to one, while REV and CON traces have lower and more disperse values. Next, we performed pairwise Kolmogorov-Smirnov tests correcting for the family-wise false discovery rate (11) (see Table S1), showing that: BOS scores are significantly higher than for REV and CON in all cases; response to SYN is significantly higher than that to REV and CON (except in ab10); and score values for SYN are significantly different than those of BOS (except in ch12).

To gain further insight into the manner in which responses to SYN differ from those to BOS, we defined “high” responses as those with score greater than 0.5 (*ρ* > 0.5). Note that close to 95% of the responses to CON and REV are below this threshold. BOS elicited a significantly larger fraction of “high” responses than SYN in 8 out of 9 experiments (Fisher’s exact test, Table S1). Interestingly, the distribution of score values for “high” responses was not significantly different between SYN and BOS (K-S test, Table S1 and Figure 4b).

The previous analysis considered the vS activity elicited by a playback motif as a whole. However, in many cases the elicited response followed the median execution pattern during only a portion of the motif (see panels ii-iii of Figure 4a). To explore the temporal pattern of activation, we identified “activity segments” as intervals of time where the measured vS activity trace closely followed the expected execution pattern of the motif (see Figure S5a and methods).

Figure 5a shows activity segments identified for playbacks of BOS and SYN for a bird. These activity segments show a structure consistent with the repetitive motif structure of the song. We therefore calculated the coverage at each time of the motif, that is, the percentage of playback motifs with detected vS activity. For this particular bird, there is a higher coverage towards the end of the motif (Figure 5b panel ii). Remarkably, the onset times of the activity segments tend to cluster at defined instants of the motif (Figure 5b panels iii-iv), producing sudden increases in the coverage percentage (indicated with arrows in Figure 5b panel ii, see also Figure S5b-e). We refer to these instants as “hotspots”.

**Figure 5.**
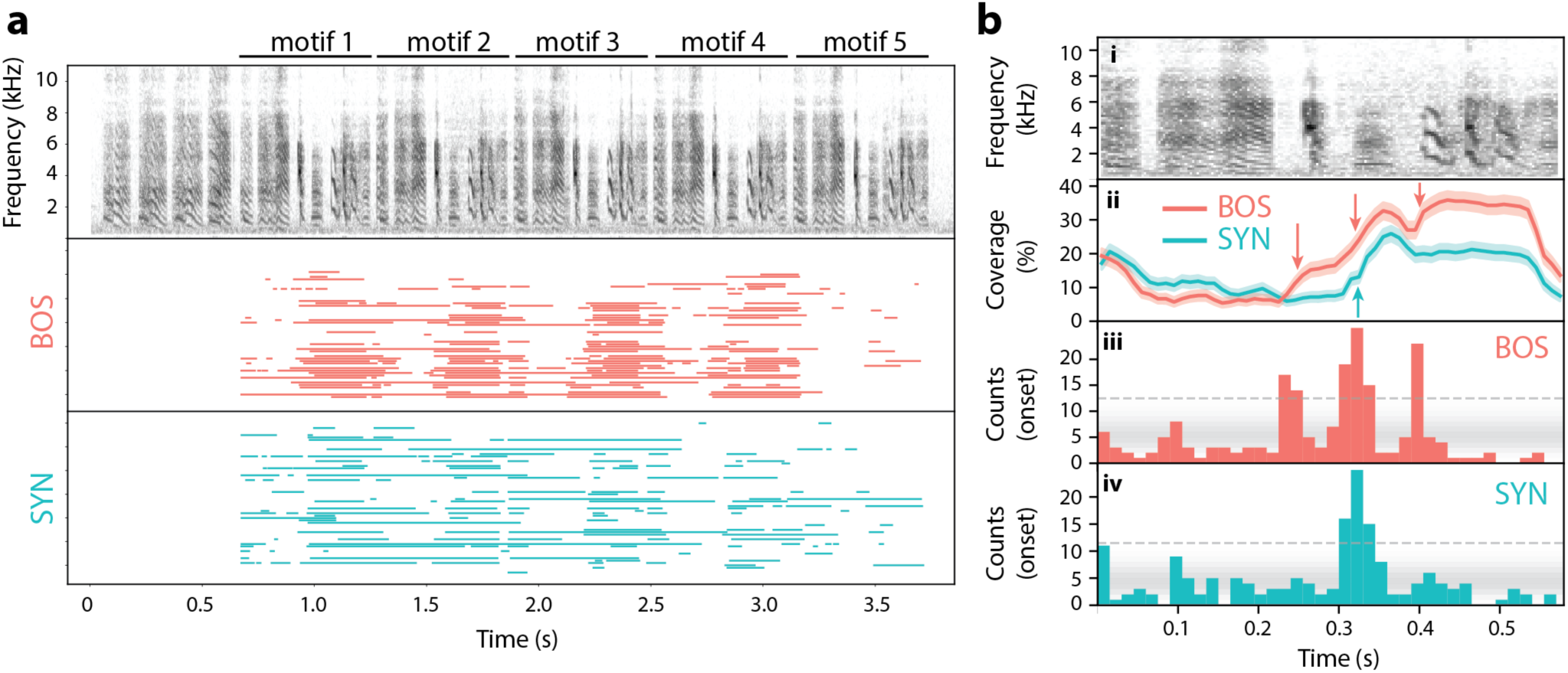
Evoked activity onsets cluster at hotspots. **a)** Top, spectrogram of full song used as auditory stimulus (bird ab05). Activity segments detected for BOS (middle, red) and SYN (bottom, blue) for all playbacks, arranged chronologically per line from top to bottom. **b)** i, spectrogram of BOS motif; ii, activity segment coverage (i.e. percentage of presented motifs that evoked vS activity) for every time of the motif, for BOS (blue) and SYN (red). Arrows indicate “hot-spots". Shaded regions correspond to the standard error as calculated by bootstrapping. iii-iv, histograms for the onset times of activity segments for BOS and SYN (15ms bins). The dashed horizontal line indicates the threshold for significant difference from a uniform distribution (using Bonferroni’s correction for family-wise error rate at α=0.05).

The coverage for the synthetic versions of the song is roughly similar to that of BOS, although significantly lower in some portions of the motif. Notably, responses to SYN show a subset of the hotspots observed for BOS (Figure 5b and S5b-e).

In some occasions, we observed vS activity patterns consistent with the execution pattern, after the last motif of the playback. Furthermore, the timing of these “virtual motifs” was consistent with what would be expected based on the inter-motif intervals (Figure 6).

**Figure 6.**
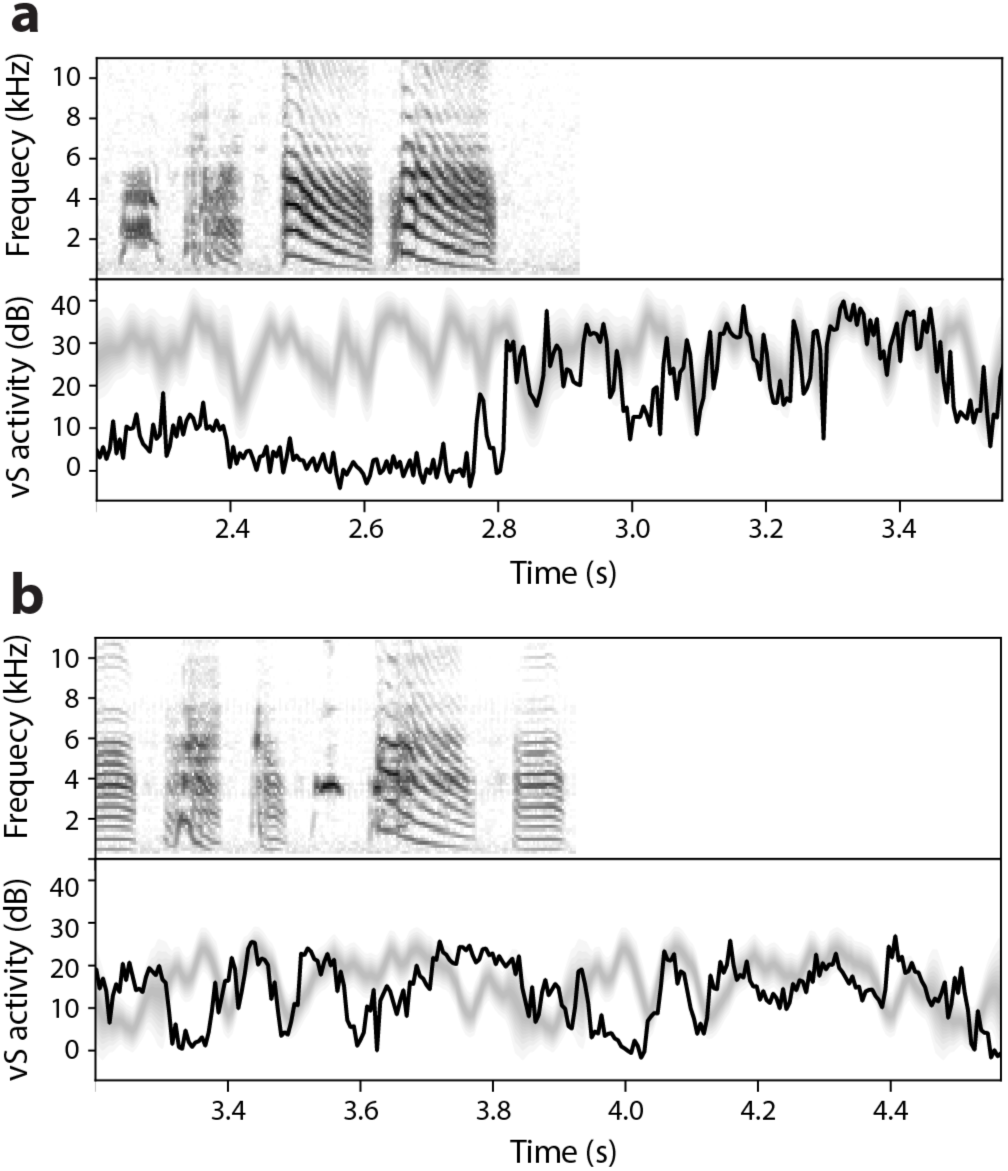
In-phase evoked activity can be detected after the end of the stimulus. **a)** Top, spectrogram of final part of auditory stimuli (ab09). Bottom, selected example of evoked vS activity pattern (black) and expected vS activity based on execution pattern and typical inter-motif intervals. **b)** Same as in **a** for bird ab11.

## Discussion

In this work, we found that nocturnal playbacks of the bird’s own song, or synthetic versions of that song, evoke vS activity patterns strikingly similar to those recorded during song execution. This result is consistent with the activity of telencephalic nuclei elicited by similar stimuli (1, 2, 5), yet it is surprising that the motor command should reach the syrinx in sleeping animals. In line with this finding, during sleep spontaneous activity patterns resembling the execution pattern can be detected in vS (but not in the respiratory gesture, explaining the lack of phonation) (8). This provides an excellent experimental opportunity to measure the integration of sensory information into a motor program.

Notably, when activated by auditory stimuli the vS signal exactly follows the execution pattern in shape and amplitude, *i.e.* it is an all-or-nothing or switch-like response (Figure 2, S2 and 3a). Note that in contrast to neuronal spikes, EMGs could potentially show partial activation in individual trials. The difference observed between autogenous and synthetic songs is the probability of eliciting a response, not the profile of the evoked response (Figure 4 and S4). Interestingly, this effect is reminiscent of the categorical perception of phonemes by humans (12, 13).

An interesting observation is that the moments at which the responses start are not uniformly distributed along the motif. Instead, they cluster at a small number of well-defined “hotspots” (Figure 5b and S5b-e). We could not correlate these hotspots to any consistent acoustic features. Our song syntheses usually produced onset of activity in a subset of the hotspots produced by BOS, which may partially explain the lower probability of evoking response by SYN.

One possible explanation for the existence of hotspots is that some acoustic features of the motif produce particularly strong auditory feedback. Alternatively, there could be specific “entry points” in the activity pattern where the dynamic tends to initiate, with no particularly strong effect of the acoustic features immediately preceding. These two interpretations are not mutually exclusive, and both could be involved in the production of the observed hotspots.

The overall delay of the vS response elicited by BOS or SYN, with respect to the execution pattern, was less than 30 ms in all cases. This is consistent with what has been previously reported for the CNS (3, 5). Furthermore, our measurements bound the delay below 10ms for several birds (not ruling out zero delay), while for others we found a near zero delay for some segments of the motif (Figure 3c-d and Table S1).

These delays are most likely shorter than the time required for the information of an auditory event to travel from the cochlea to the CNS (6 synapses), be processed by the song system (at least 2 synapses) and then relayed to the syrinx (Figure 1a) (14, 15). Therefore, our data suggests that a specific phonologic event can only evoke activity patterns that come later in the motif, not the gesture causally related to the production of that particular event. In this sense the song system appears to be working in a predictive manner (16). Supporting this idea, in some infrequent cases vS activity continued after the end of the last motif, in a pattern consistent with what would be expected for an additional “virtual” motif (Figure 6).

In order to function in a predictive manner, the song system has to have some sort of preprogrammed internal dynamics. The correct sequence of auditory stimuli could then entrain this dynamic, activating the system and evoking the response. This conceptual model is consistent with the differences observed between BOS and SYN; a suboptimal sequence of auditory stimuli, as the one produced by our synthetic song, is less efficient at inducing the response, but once induced the response follows its preprogrammed dynamic, independently of the stimulus that evoked it (Figure 2, s2 and 4). Furthermore, this model is also consistent with the variable time delays observed in some birds (Figure 3d), as coupling of non-linear dynamical systems ordinarily results in phase differences between systems that evolve in time (17).

In this way, the study of a motor output elicited by synthetic stimuli, revealed the predictive and switch-like nature of a sensory-motor integration program.

## Methods

### Subjects

Adult male zebra finches (*Taeniopygia guttata*) were acquired from local breeders. After a 10 days quarantine protocol, birds were housed in large cages (40×40×120 cm) with conspecific males under a 14h/10h light/dark cycle, had *ad libitum* access to food and water, and could interact visually and vocally with a conspecific female. Experimental protocols were approved by the Animal Care and Use Committee of the University of Buenos Aires.

### Song recording

Bird’s songs were recorded 3-5d prior to the surgery in a custom-built acoustic isolation chambers with a microphone (Takstar SGC568) connected to an audio amplifier and a data acquisition device (DAQ, National Instruments USB-6212) connected to a PC. Custom Matlab v7.10.0 (The Mathworks) scripts were used to detect and record sounds, including a 1s pre-trigger window.

### Surgery

Custom made bipolar electrodes were implanted in the vS muscle as previously described (6) and connected to an analog differential amplifier (225x) mounted on a backpack previously fitted to the bird (see details in SI Methods).

### EMG recording

After a 4h recovery time, birds were place in the acoustic isolation chamber and the amplifier mounted to the backpack connected to the DAQ through a rotator to allow free movement. Two external 12V batteries powered the amplifier in order to avoid line noise. EMG and audio signals were recorded by a PC running Matlab, at a sampling rate of 44150Hz. During day/night recordings were triggered by sound/EMG with a 1s pre-trigger window. We adjusted trigger thresholds for each bird to levels slightly higher than the basal range, minimizing the probability of missing events, even when this led to a high percentage of spurious records.

### Playback stimuli

A synthetic version (SYN) of the bird’s own song (BOS) was produced as previously reported (18) (see details in SI Methods). Playback protocols were passed at night (11-12pm to 5-6am) from a speaker inside the acoustic isolation chamber. Each protocol consisted of several stimuli of similar duration including BOS, SYN, reverse BOS (REV) and a conspecific song (CON), presented in random order. 15s were recorded per playback, starting 3s before the stimulus onset. Protocols were separated by a random 10.0±0.4min interval of silence. Playback volume was set at 55±5dB (calibrated with 4kHz beeps, YF-20 Sound Level Meter, Yu Fung).

### Data analysis

Recordings were analyzed in Pyhton v3.5.2 using custom scripts (available at github.com/Dynamical-Systems-Lab). See details in SI Methods. The envelope of the EMG signal was calculated in the following manner. First, the Teager energy operator was applied (10), defined as

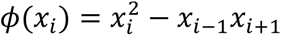

Then the 95^th^ percentile for each 5ms bins (220 samples) was determined, resulting in *f*_*s*_ = 200*Hz* vector with a robust estimation of the EMG’s Teager energy envelope (*rϕ*). We found that this method greatly attenuated the electrocardiogram component of the signal observed in some birds (Figures S1a). We then defined

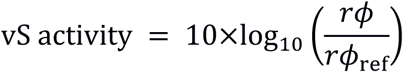

where *rϕ*_ref_ is the mode of *rϕ* during periods of no activity. Note that vS activity is measured in dB.

To compare the elicited activation patterns independently of the variable delays observed (Figure 3d) we implemented a smooth time warping (STW) algorithm that allows small nonlinearities in the time stretching (Figure S3c, see details in SI Methods).

To quantify the response elicited by the different stimuli, we defined the normalized projection score *ρ* as

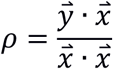

where 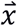 is the median of the execution patterns (EXE_50_) and 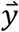 is the vS activity trace for a particular playback trial or execution pattern. Before calculating the score, we time-aligned all the vS activities to the average execution pattern of a selected song using our STW algorithm.

### Statistical analysis

Using the response score *ρ*, we performed all pairwise comparisons between stimuli for each bird using a Kolmogorov-Smirnov (KS) test and controlled for false discovery rate at *α* = 0.05 (11). Results are informed with letter code in Figure 4b. We compared the proportion of high responses (*ρ* > 0.5) between BOS and SYN using Fisher’s exact test for count data. Finally, we compared the distribution of the high responses between BOS and SYN using a KS test. P-values for these tests are informed in Table S1. All statistical tests were done using R v3.4.2 (19).

Periods of evoked vS activity consistent with the execution pattern were detected in the following manner (see Figure S5a). The baseline distribution of vS activity *P*_0_(*vS*) was estimated from −3 to −1 seconds relative to playback onset. The time dependent distribution of vS activity during motif executions *P*_*x*_(*vS, t*) was estimated from data as that of Figure 1c, time-locked to each playback motif. These were then used to calculate the log-probability ratio defined as

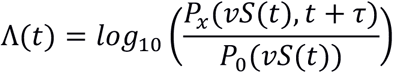

where *vS*(*t*) is the measured vS activity at time t and *τ* is the average time delay between the evoked response and the execution pattern as calculated in Table S1. Segments of evoked activity between times *t*_*i*_ and *t*_*f*_ were identified according to the following conditions:

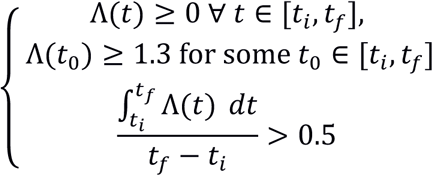

Segments of less than 15ms of duration were removed and consecutive segments separated by small gaps were consolidated if the gap accounted for less than 10% of the resulting segment. This algorithm was benchmarked with manual classification of activity segments giving qualitatively similar results.

## Acknowledgements

We would like to thank M. A. Suarez for veterinarian support, S. Boari, A. Sanchez, R. G. Alonso, J. Lassa Ortiz and C. T. Herbert for help with animal care, and A. Amador for useful suggestions and discussion. This work was supported by the National Council of Scientific and Technical Research (CONICET, Argentina), the National Agency of Science and Technology (ANPCyT, Argentina), the University of Buenos Aires (UBA) and the National Institute of Health (NIH) through R01-DC-012859 and R01-DC-006876.

## Author contributions

G.B.M. and F.G. conceived the project, A.B. and G.B.M. designed research, A.B. performed researched and analyzed data, J.F.D. developed the EMG recording electronics, A.B. and G.B.M. wrote the paper.

The authors declare no conflict of interest.

## Supporting Information

### SI Methods

#### Electrodes

Bipolar electrodes were prepared manually using ultrafine wire (Ø 25μm, stainless steel 304, heavy polyimide HML insulated, annealed, California Fine Wires Company, item #CFW0013157). The insulation-peeled tips were twisted onto themselves (~3 turns/mm) to increase rigidity and facilitate implantation. A Ø~0.3mm drop of epoxy (Cytec Easypoxy K-20) was placed ~0.5mm from the tip to improve attachment to the muscle. A reference electrode of multifilament copper fine wire (Ø 90μm, Phoenix Wire Inc.) was used, which also served as mechanical scaffold for the vS electrodes facilitating manipulation and increasing mechanical resistance.

#### Surgery

1-2d before surgery we fitted a Velcro backpack to each bird and attached a 1.6g weight for habituation. Prophylactic antibiotic was administered orally (chloramphenicol ~25mg/day, Cloralen) starting one day before the surgery. Food and water were removed 1h prior to the surgery and 10m before surgery 0.5mg/Kg of midazolam benzodiazepine (Dozilam) was administered orally.

Animals were anesthetized with isoflurane (Forane, Ohio vaporizer model 100F) administered through a custom build gas mask at an airflow of 1.6 L/min and concentrations in the range 0.75-1.5 % v/v, adjusted as required. Anesthetic depth was assessed based on respiration rate and lack of response to gentle pinches on the skin and feet. Lidocaine gel (2%, Veinfar) was used as topical anesthetic and to separate the feathers.

Using forceps (Dumont #5 and #7 biology tip), chirurgical scissors (Vannas straight 8mm blades, Reda) and a surgical microscope (Zeiss OPMI 1), we made a ~1cm incision rostral to the keel and removed or displaced adipose tissue as required to expose the interclavicular air sac. After removing any fluid from the air sac surface using paper points (Meta biomed), we made a ~1-2mm incision on the air sac, ripped the fascia and implanted two electrodes on the exposed *vS* muscle using a needle holder (Fine Science Tools). We fixed the electrodes using 1-3 drops (~1ul) of 2-octyl cyanoacrylate tissue adhesive (Surgi-lock 2oc, Meridian) delivered by a custom build precision dispenser.

Next, we reconstructed the air sac using adipose tissue and sealed it with cyanoacrylate adhesive. We passed vS and reference electrodes subcutaneously towards the back where they exited dorsally through a hole in the Velcro backpack. We then sutured the ventral incision (Ethilon 6-0 SC-16) and released the animal from anesthesia.

Finally, we cut the electrodes at adequate length, welded them to pin headers and connected them to a custom-built circuit attached to the backpack. This circuit consisted of a low power differential amplifier (AD620, Analog Devices) set at 225x gain and a passive high-pass RC filter with a 150Hz cutoff frequency, with an overall weight of 1.6g. We used hot-melt adhesive to increase mechanical resistance of the electrode cables and to place them in a position where the bird could not peck at them.

#### Song synthesis

The following system of equations was integrated using the 4^th^ order Runge-Kutta method implemented with the package Numba v0.29.0 for Python.

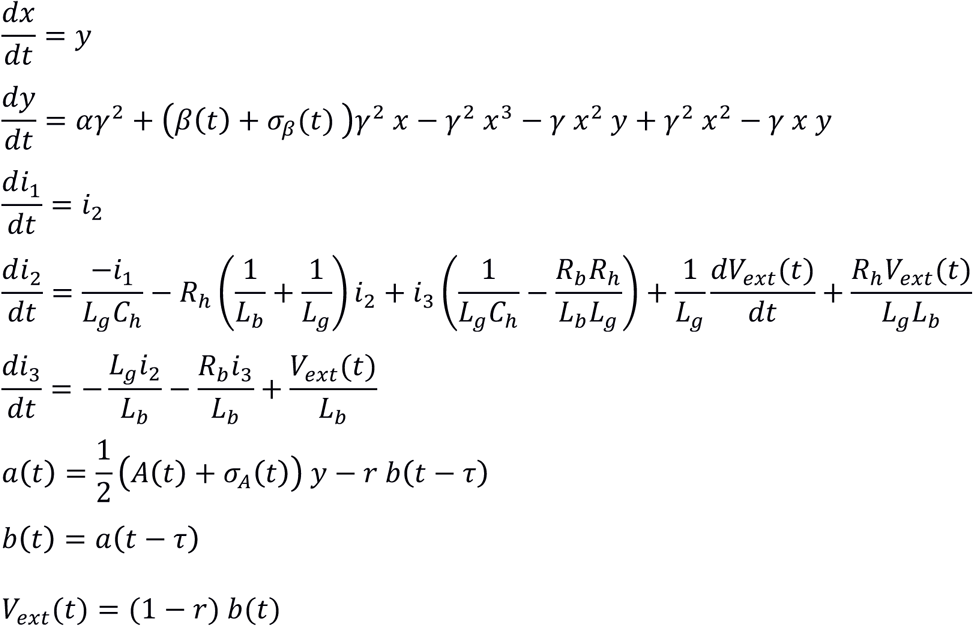

The synthesis was tuned for each bird adjusting parameters in the following ranges: *r* ∈ [0.40,0.61], *L*_*g*_ ∈ [28,140], *L*_*b*_ = 209, *R*_*b*_ = 1.04×10^7^, *R*_*h*_ ∈ [0.1,2.0]×10^5^, *τ* ∈ [1.7,2.3]×10^−4^, *C*_*h*_ ∈ [3.2,7.8]×10^−11^, *γ* = 2.4×10^4^ and *α* = −0.15. The time-dependent amplitude parameter *A*(*t*) was set to the envelope of the BOS oscillogram, normalized so that the 99% percentile had a value of 10 (*i.e.* a normalization robust to outlier samples). A uniform random noise *σ*_*A*_(*t*) of amplitude in [0,0.05] was added to *A*(*t*). The time dependent parameter *β*(*t*) was calculated from the fundamental frequency *ℱ*(*t*) according to (1), which can be approximated by the following equation:

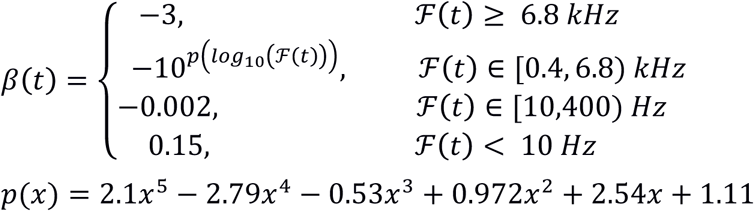

*ℱ*(*t*) was manually fitted to the BOS spectrogram using praat v6.0.36 (2). A normal random noise *σ*_*β*_(*t*) with standard deviation in [0, 0.02] was added to *β*(*t*). The output was defined as sound = *R*_*b*_*i*_3_.

#### Data Analysis

Sound/EMG files were pre-processed using a 4^th^ order Butterworth highpass/bandpass filter with cutoff frequencies of 300Hz/300-3000Hz. Abnormal EMG recordings, in which the signal was flat, saturated or followed the audio signal were removed. These artifacts normally preceded electrode detachment.

Time intervals of sound were detected by thresholding the envelope of the audio oscillogram. Within these intervals, we searched for instances of the motif execution by correlating the spectrogram (12ms Hanning window, 6ms overlap) to the spectrogram of a manually selected motif template. To correctly align motif executions and compensate for small differences in motif duration, we calculated a linear time warping on the audio signal and applied this transformation to the vS recording.

To determine the activation pattern elicited by the different stimuli, vS activities traces were time-locked to the presented motifs and the 95^th^ percentile at each time point was calculated (referred to as BOS_95_, SYN_95_, CON_95_ and REV_95_). These patterns were correlated with the median vS activity of the motif during song execution (EXE_50_), according to the following expression

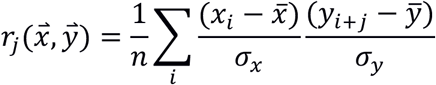

where *r*_*j*_ is the correlation at a delay of *j* samples (*τ* = *j*/*f*_*s*_), 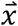 is the reference patter (EXE_50_) with mean 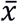 and standard deviation *σ*_*x*_, and 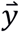 is the test pattern (*e.g.* BOS_95_, SYN_95_, etc.).

We constructed “correlation spectrograms” (Figure 3d bottom) by taking 250ms time windows of the evoked vS activity and calculating the local Pearson correlation (*r*_*j*_) with the execution pattern, for different delays. Then we took partially overlapping time windows and repeated until the duration of the playback was covered.

Multidimensional scaling (MDS) was performed with Python’s package scikit-learn v0.19.1. vS activity traces were time-aligned to the average execution pattern of a selected song using our STW algorithm (see below) and Euclidean distances between the resulting traces were calculated. The positions of the MDS points, which correspond to individual responses, were initialized using the first two components of a principal component analysis (PCA). In this way, the starting positions of the strain minimization procedure were comparable across animals.

#### Smooth Time Warping

First, we calculated a smooth interpolation function *f*(*t*) such that *f*(*t*_*i*_) = *y*_*i*_ for all time points *t*_*i*_, where *y*_*i*_ is the corresponding vS activity. Next, we defined the time-warping function *ω*(*t*) and the time warped vector 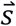 such that *s*_*i*_ = *f*(*ω*(*t*_*i*_)). We then searched for the time-warping function that maximizes the correlation between 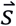 and the reference pattern 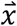, that is 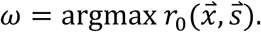 To this end we defined the delay function *δ*(*t*) = *ω*(*t*) — *t* and parameterized it with a vector of M parameters 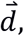 such that *δ*(*τ*_*i*_) = *d*_*i*_, with evenly spaced interpolation times *τ*_*i*_. This parameterization has the advantage that it results in a fixed density of parameters per unit time, and therefore performance does not depend on the length of 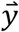 Another advantage is that the parameters are easily interpretable as the delays in seconds with the reference pattern. We did the optimization by minimizing the regularized cost function

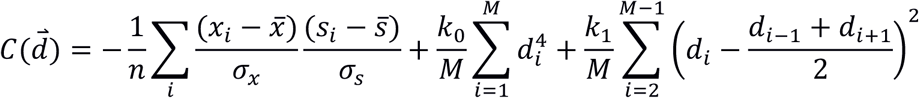

where the first term reflects the cost reduction as the correlation increases and the other two terms penalize for large delays and departures from linearity, respectively. We used *k*_0_ = 1.6×10^5^, *k*_1_ = 10^4^, Δ*τ* = *τ*_*i*_ — *τ*_*i*−1_ = 0.1s, interpolated using cubic splines as implemented in NumPy v1.13.0 (3) and used SciPy v1.0.0 (4) simplex optimization algorithm.

## SI Figures

**Figure S1.**
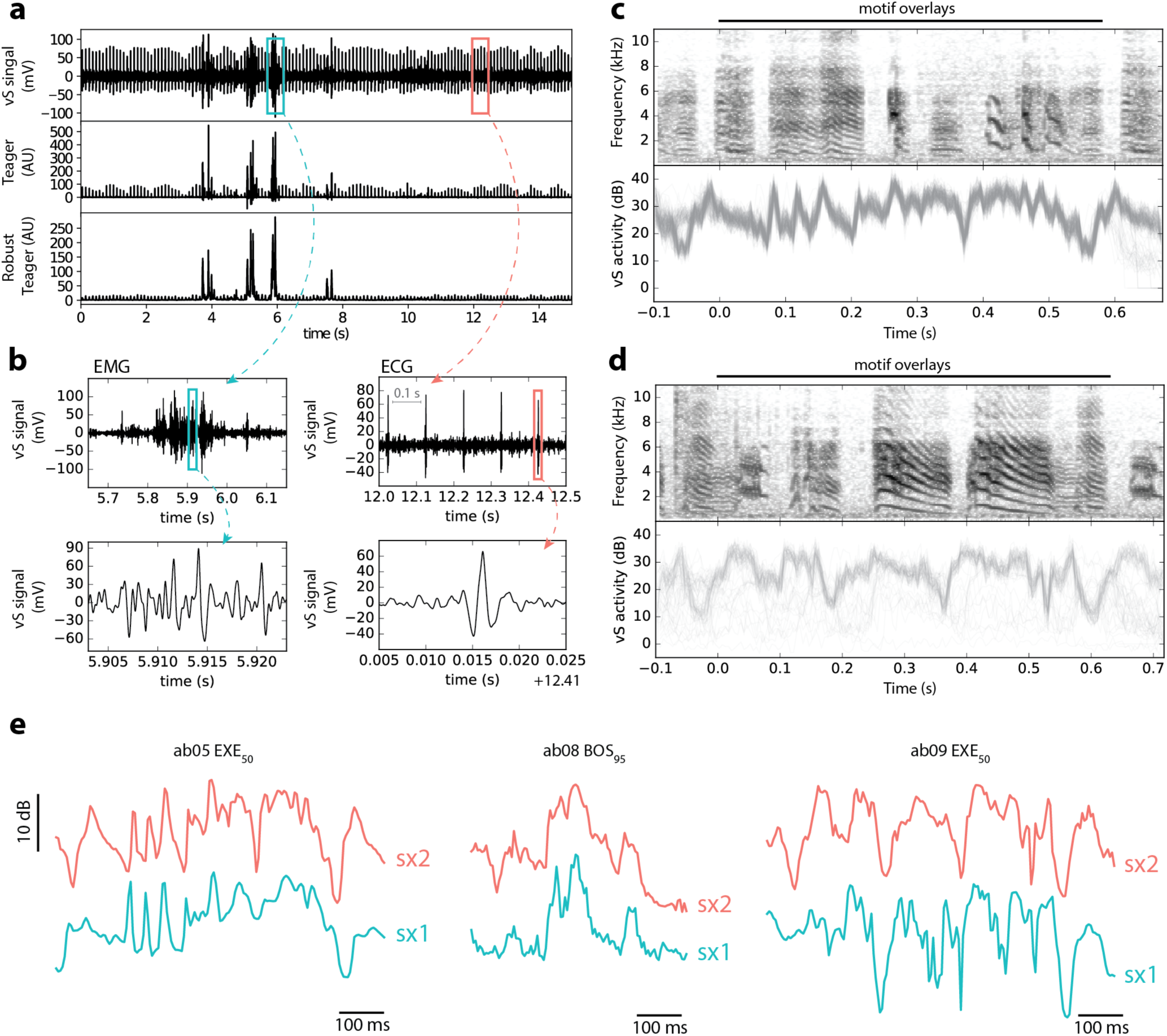
vS envelope method and execution patterns. **a)** Top: raw signal of vS after amplification. Middle: envelope calculated applying the Teager energy operator. Bottom: robust estimation of the maximum Teager energy in 5ms bins (see methods). **b)** Zoom-in of indicated regions of the raw signal. **c)** Top: Spectrogram of the motif of bird ab05. Bottom: overlay of all vS activity execution traces. **d)** Same as **c** for second surgery of ab09 (compare with Figure 1c). **e)** Comparison of average vS activity pattern between the measurements obtained in two independent implantation surgeries (Sx1, blue; Sx2, red). For ab05 and ab09 the median execution profile is shown (EXE_50_). Bird ab08 did not sing while implanted, so the 95^th^ percentile of the response to the bird’s own song playback (BOS_95_) was used instead.

**Figure S2.**
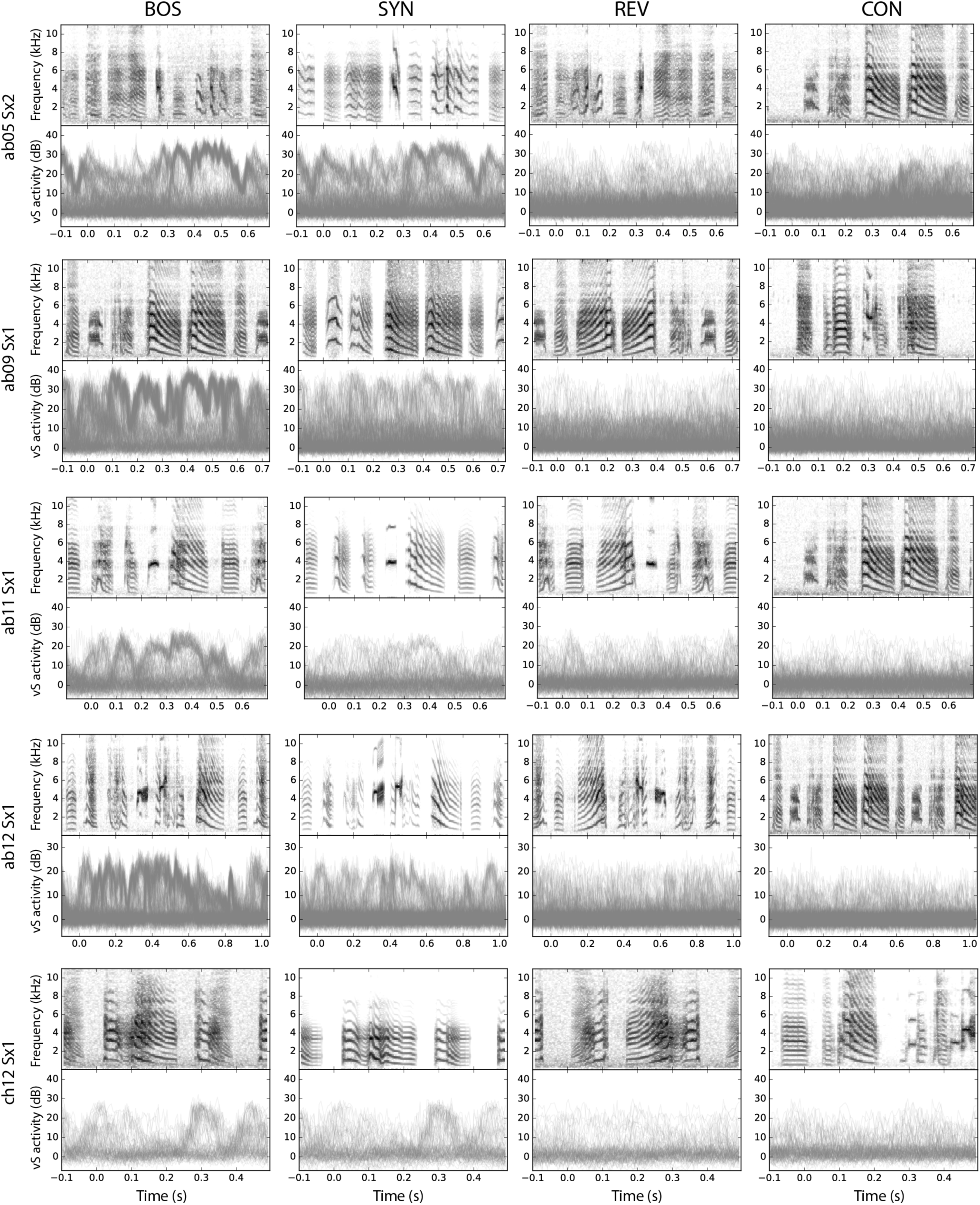
Response to nocturnal playbacks. Top of each panel: spectrogram of the stimulus motif. Bottom of each panel: overlay of all vS activity traces measured during playbacks of that motif. Each column corresponds to the indicated stimulus (BOS, SYN, REV and CON) and each row corresponds to the indicated bird/surgery.

**Figure S3.**
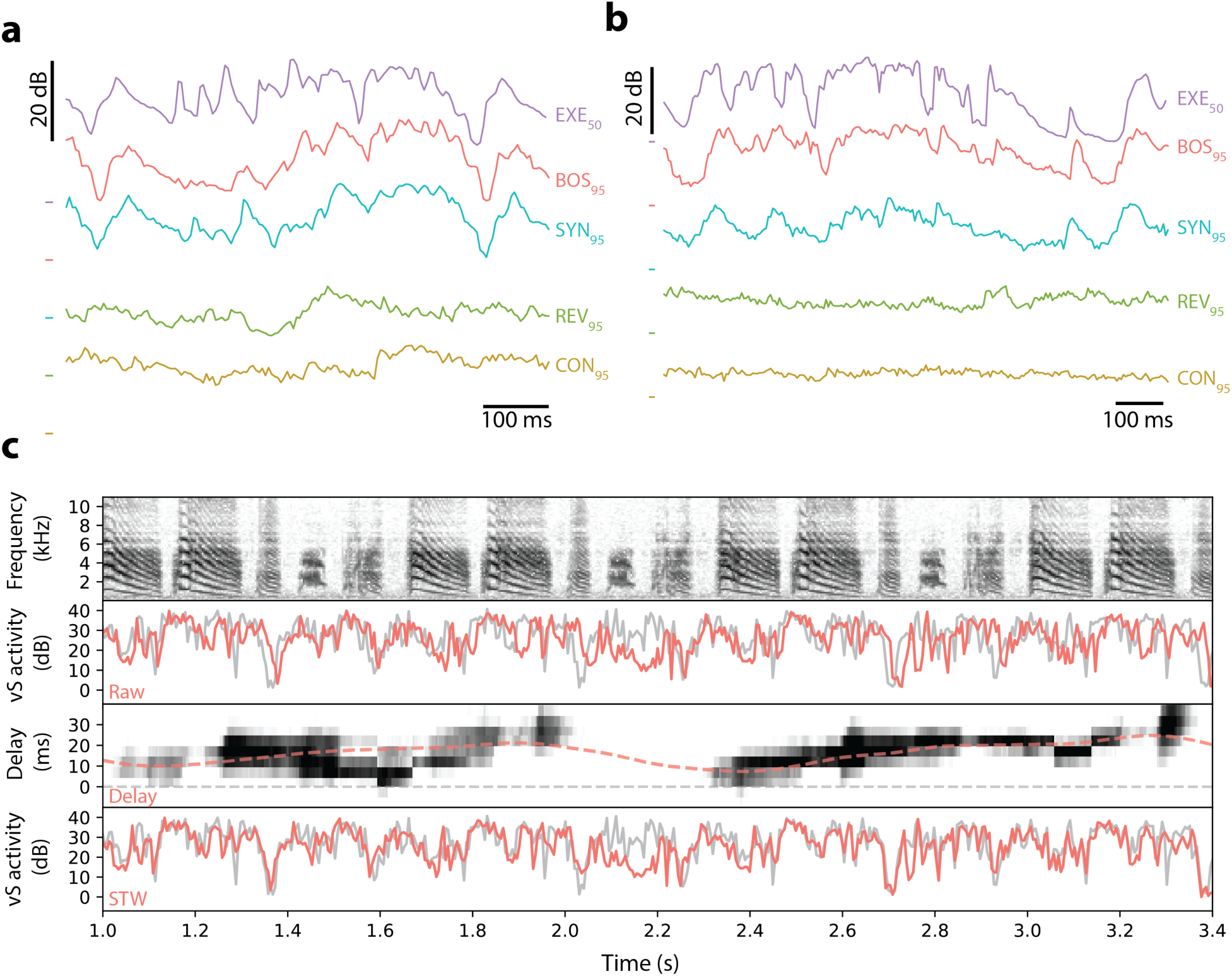
Evoked vS activity patterns and Smooth Time Warping algorithm. **a)** vS activity median execution pattern (EXE_50_) and 95^th^ percentile of the responses evoked by nocturnal playbacks (BOS_95_, SYN_95_, REV_95_ and CON_95_) for bird ab05. **b)** same as **a** for bird ab12. **c)** Top panel: spectrogram of song of ab09 used as playback stimulus. Second panel: vS activity during the execution of that song (gray) and evoked during an auditory stimulus (red). Third panel: correlation spectrogram showing the local correlation coefficient (in gray scale) between evoked and execution traces, for every time and delay. The red dashed curve shows the delays calculated by the Smooth Time Warp algorithm (STW, see SI Methods). Bottom: execution vS activity trace (gray) and evoked response after applying smooth time warping (red).

**Figure S4.**
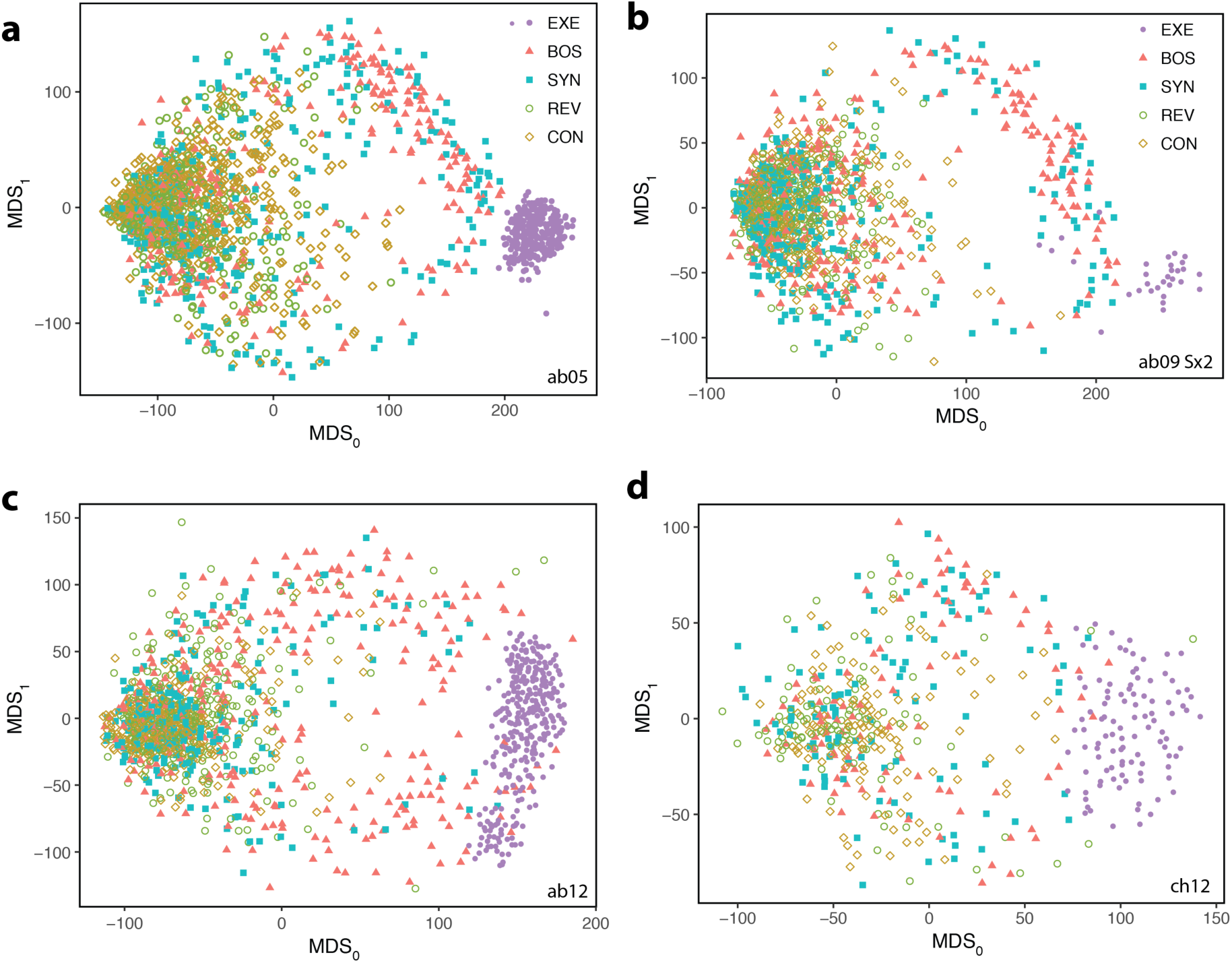
Multi-Dimensional Scaling of vS activity execution traces (EXE, violet) and responses to nocturnal auditory stimuli (BOS, red; SYN, blue; REV, green; CON, brown). **a)** bird ab05. **b)** bird ab09, second surgery. **c)** bird ab12. **d)** bird ch12.

**Figure S5.**
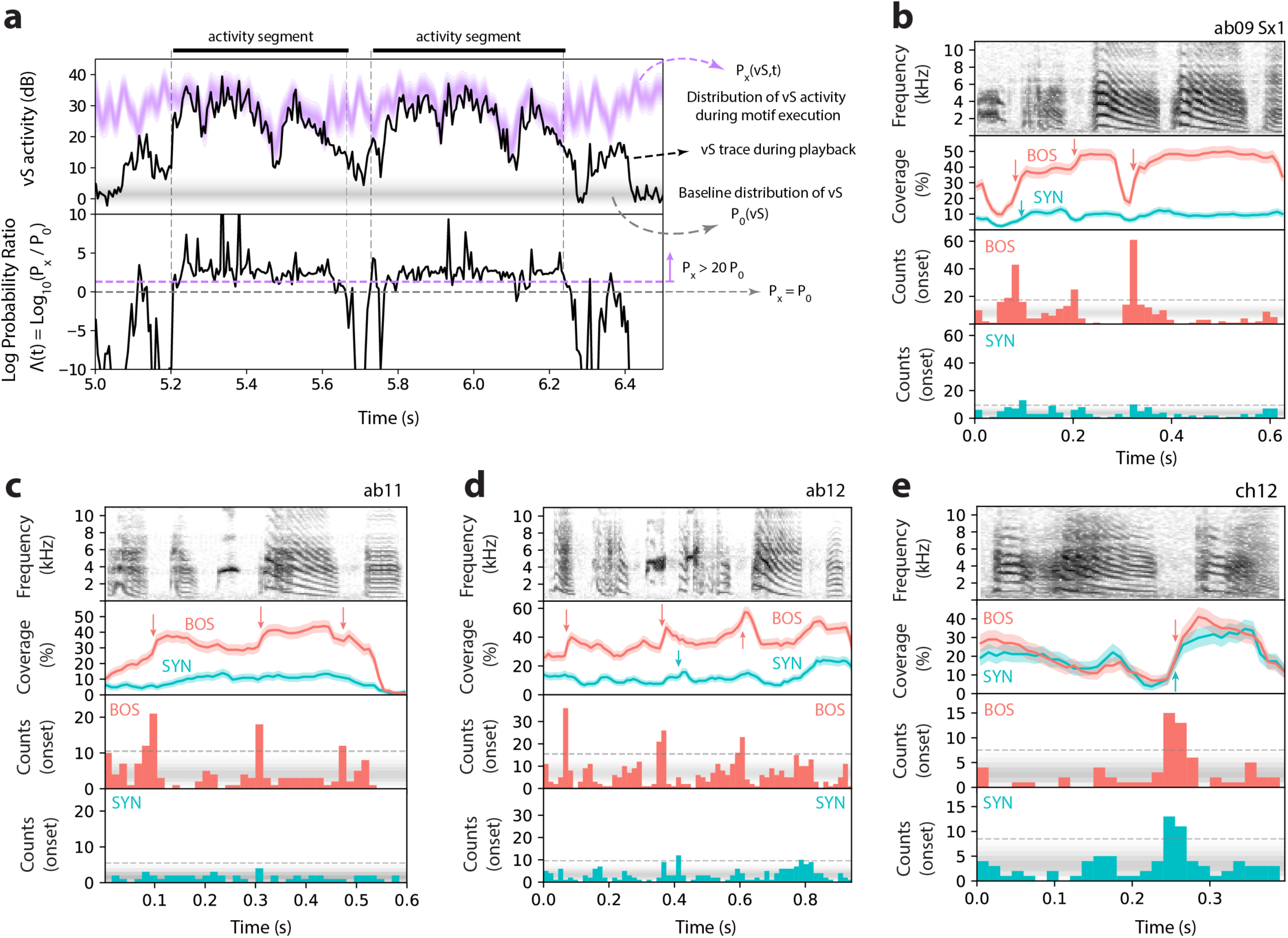
Detection of vS activity segments and hotspots. **a)** Example of activity segment detection algorithm. Top: The vS activity trace evoked during a particular playback is shown in black. The violet shaded region denotes the expected vS activity distribution as calculated from the execution profiles *P*_*x*_(*vS, t*), and the gray shaded region the distribution of baseline vS activity *P*_0_(*vS*). Bottom: Log Probability Ratio evolution calculated as Λ(*t*) = *Log*_10_(*P*_*x*_(*vS*(*t*),*t*)/*P*_0_(*vS*(*t*))). The horizontal dashed lines correspond to the thresholds defined by *P*_*x*_ = *P*_0_ (gray) and *P*_*x*_ = 20 *P*_0_ (violet). “Activity segments” are shown with horizontal bars over the plot. See methods for details. **b)** Spectrogram of playback motif and activity segment coverage for BOS (blue) and SYN (red), for bird ab09 Sx1. The shaded regions represent the standard error of each curve as calculated by bootstrapping. Bottom panels show activity onset histograms for BOS and SYN, showing the significance threshold as gray dashed line (assuming a uniform distribution and controlling for family-wise error rate). **c)** same as **b** for bird ab11. **d)** same as **b** for bird ab12. **e)** same as **b** for bird ch12.

## SI Tables

**Table S1.** Delays, correlation coefficients and statistics. **Bird:** animal identifier. **Sx:** surgery number. **type:** playback stimulus type. **n:** number of trials. **τ±err:** delay of the correlation between the 95% percentile of all the responses to this stimulus and the median execution pattern (n.d.: not determined). The error τ was calculated as a combination of the standard error as calculated by bootstrapping and the instrumental error (Teager envelopes were sampled at 200Hz). **cor±se:** correlation coefficient and standard error calculated by bootstrapping. **p K-S:** p-value of Kolmogorov-Smirnov test between all the points in BOS and SYN. **Rsp^h^:** number of “high” responses above the score threshold of 0.5. **Pct^h^:** percent of high responses (Rsp^h^/n). **p Fisher^h^:** p-value of Fisher’s Exact Test for Count Data, for high/low responses to BOS/SYN. **p K-S^h^:** p-value of Kolmogorov-Smirnov test between BOS and SYN, considering only points with high scores (*ρ* >0.5). † the marginally significant difference for ab05 vanishes if a threshold of *ρ* > 0.49 is used for “high” values. **score^h^±se:** mean value of the score for high responses (score > 0.5). Standard error of the mean was calculated by bootstrapping. **sd^h^±se:** standard deviation of the score for high responses (score > 0.5). Standard error of the standard deviation was calculated by bootstrapping.

| bird | Sx | type | n | $\tau \pm \text{err}$<br>(ms) | corr±se | p K-S | Rsp<sup>h</sup> | Pct<sup>h</sup> | p Fisher<sup>h</sup> | p K-S<sup>h</sup> | score<sup>h</sup>±se | sd<sup>h</sup>±se |
| --- | --- | --- | --- | --- | --- | --- | --- | --- | --- | --- | --- | --- |
| ab05 | 1 | bos | 640 | 0±5 | 0.75±0.02 | | 57 | 9% | | | 0.60±0.01 | 0.068±0.007 |
|  |  | syn | 0 |  |  |  |  |  |  |  |  |  |
| | 2 | bos | 315 | 14±5 | 0.72±0.02 | 0.0013 | 107 | 34% | 0.000034 | 0.043<sup>†</sup> | 0.68±0.01 | 0.114±0.005 |
| | | syn | 320 | 16±5 | 0.63±0.02 | | 62 | 19% | | | 0.71±0.02 | 0.128±0.007 |
| ab08 | 1 | bos | 85 | | | 0.00028 | 26 | 31% | 0.0015 | 0.67 | 0.68±0.03 | 0.14±0.02 |
| | | syn | 72 | | | | 7 | 10% | | | 0.67±0.04 | 0.11±0.04 |
| | 2 | bos | 145 | | | <1e-6 | 82 | 57% | <1e-6 | 0.81 | 0.64±0.01 | 0.092±0.008 |
| | | syn | 116 | | | | 14 | 12% | | | 0.66±0.03 | 0.13±0.02 |
| ab09 | 1 | bos | 296 | 8±5 | 0.67±0.02 | <1e-6 | 146 | 49% | <1e-6 | 0.13 | 0.78±0.01 | 0.112±0.006 |
| | | syn | 296 | 18±5 | 0.65±0.03 | | 32 | 11% | | | 0.74±0.02 | 0.13±0.01 |
| | 2 | bos | 288 | 17±5 | 0.69±0.03 | <1e-6 | 72 | 25% | 0.0010 | 0.16 | 0.68±0.01 | 0.104±0.007 |
| | | syn | 288 | 11±5 | 0.77±0.03 | | 40 | 14% | | | 0.72±0.02 | 0.119±0.009 |
| ab10 | 1 | bos | 41 | 1±7 | 0.64±0.09 | <1e-6 | 9 | 22% | 0.00076 | | 0.68±0.03 | 0.08±0.02 |
| | | syn | 45 | n.d. | 0.38±0.04 | | 0 | 0% | | | | |
| ab11 | 1 | bos | 198 | 26±5 | 0.83±0.01 | <1e-6 | 61 | 31% | <1e-6 | 0.83 | 0.68±0.01 | 0.11±0.01 |
| | | syn | 165 | 25±15 | 0.74±0.06 | | 14 | 8% | | | 0.68±0.03 | 0.12±0.02 |
| ab12 | 1 | bos | 295 | 13±5 | 0.75±0.02 | <1e-6 | 98 | 33% | <1e-6 | 0.35 | 0.73±0.01 | 0.144±0.008 |
| | | syn | 236 | 0±5 | 0.62±0.05 | | 20 | 8% | | | 0.68±0.03 | 0.13±0.01 |
| ch12 | 1 | bos | 100 | 3±22 | 0.77±0.04 | 0.64 | 18 | 18% | 0.10 | 0.27 | 0.63±0.02 | 0.08±0.01 |
| | | syn | 104 | 3±5 | 0.76±0.06 | | 10 | 10% | | | 0.66±0.02 | 0.06±0.01 |

